# Efficient Double Helix Detection with Steerable Filters

**DOI:** 10.1101/2025.08.14.670427

**Authors:** Andrew E. S. Barentine, Ashwin Balaji, W.E. Moerner

## Abstract

We present an efficient detection scheme for localization of Double Helix point-spread functions for 3D single-molecule localization microscopy or tracking. Using steerable filters, we extract both 2D position and lobe orientation (axial position) estimates using just 7 convolutions, orders of magnitude less than used in deep-learning-based approaches. We pair this detection with a fitter and implement both as a plug-in for the open source PYthon Microscopy Environment (PYME), which features percentile-based background subtraction, signal-to-noise-based detection thresholding, and performant parallel analysis. Our complete localization analysis pipeline achieves state-of-the-art performance with minimal user input.

## Introduction

Point-spread functions (PSFs) engineered to encode additional information for single-molecule localization microscopy (SMLM) can become considerably more difficult, and computationally inefficient, to detect and localize. Open-aperture PSFs can be readily detected on a camera frame using a matched difference of Gaussians filter, which requires just 2 (separable) filterings of the camera frame [1]. However, engineered PSFs can produce far more complex spatial patterns that may or may not change significantly with defocus. The requirement to detect emitters on many (10^4^ - 10^7^) camera frames makes computational speed an imperative, and the accuracy of initial parameter estimates can additionally improve the performance of localization fitting.

The Double Helix (DH) is an engineered PSF with two lobes which rotate with defocus, and as such is typically used to encode the axial position of emitters in the orientation of the two lobes of the imaged PSFs [2, 3] as shown in Figure 1A. The DH PSF is often modeled using a double Gaussian function,

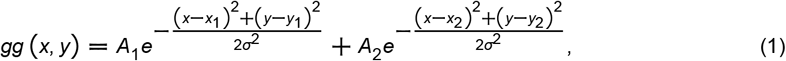

where *A*_*i*_, *x*_*i*_, and *y*_*i*_ are the amplitudes and center positions of each Gaussian, and *σ* is the shared width parameter. For the DH PSF, we find it more natural to perform a change of variables such that the center position of the PSF, (*x*_∘_, *y*_∘_), the separation between its lobes, *L*, and lobe orientation, *θ*, (see Fig. 1B) can be directly fitted:

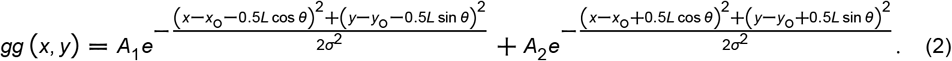

**Figure 1:**
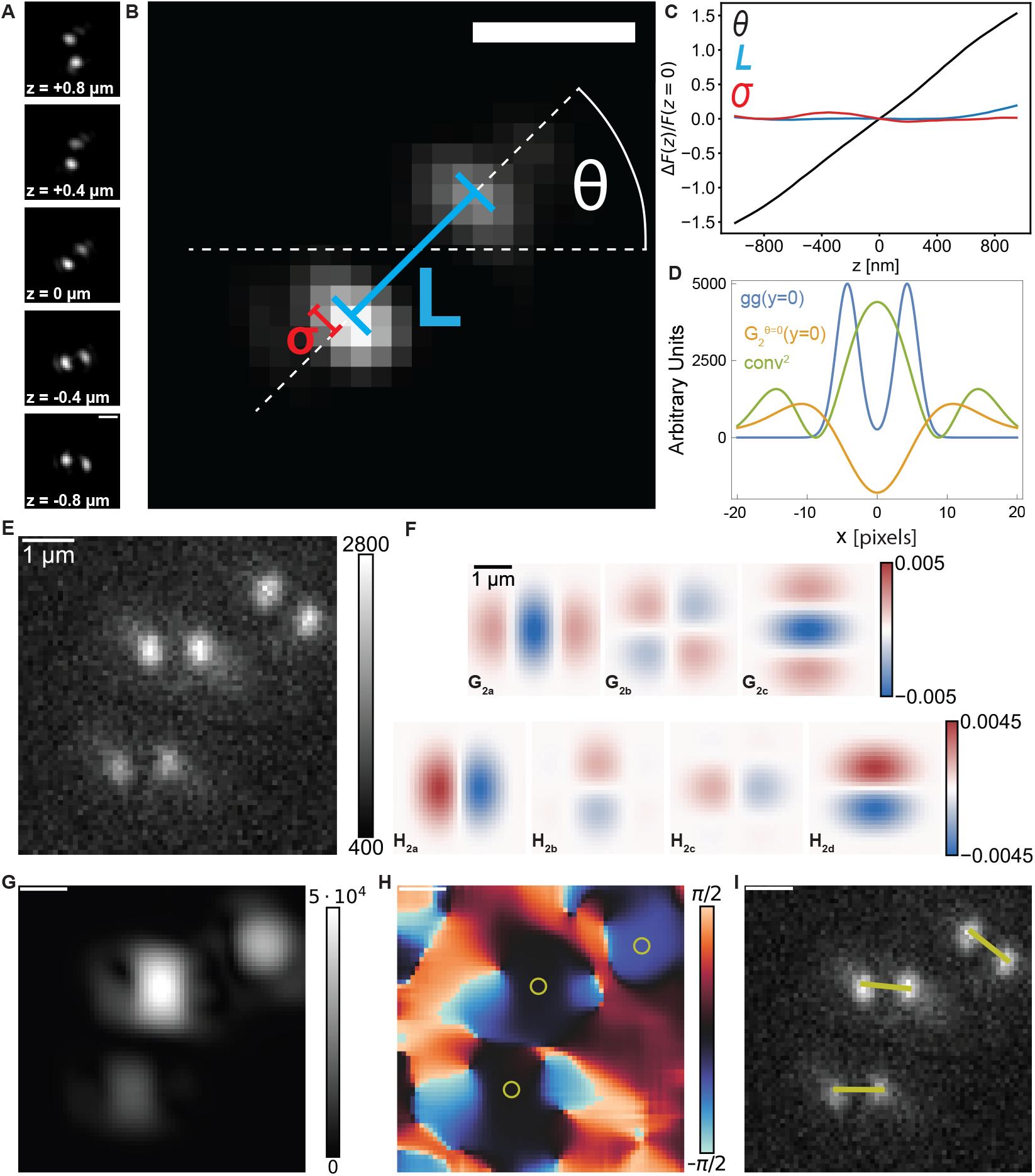
A) An experimental DH PSF z-stack. B) The z = 0 nm slice from (A), showing the lobe width parameter *σ*, the lobe separation parameter *L*, and the orientation parameter *θ* for the double Gaussian model of eq. 2. C) Fitted calibration curves for *θ, L*, and *σ*, all shown as fractional changes relative to their z = 0 values. D) 1D squared convolution of the double Gaussian model *gg* (*y* = 0), with parameters *A*_1_ = *A*_2_ = 5000 and *L* = 8.54, *σ* =1.58, *σ*_*f*_ = 6.25 [pixels].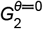 (*y* = 0) is plotted on its own scale. E) A simulated DH PSF SMLM frame from [12]. F)The separable *G*_2_ and *H*_2_ basis filter kernels for *σ*_*f*_ = 5.5 [pixels]. G) The strength map resulting from filtering (E). H) The orientation map resulting from filtering (E). Yellow circles highlight detected emitter positions. I) Detected emitter positions (maxima of the strength map) and orientations drawn as yellow sticks overlaid on (E). All scale bars are 1 µm.

With defocus across the designed axial range, *θ* rotates roughly *π* rad, yet the overall double-lobed structure of the DH PSF remains consistent, with minimal changes to *L* and *σ* (see figure 1B,C). In addition to efficient detection of candidate molecules for fitting, the ideal detection algorithm would provide accurate estimates for (*x*_∘_, *y*_∘_) and *θ* for each candidate.

The classic method of detection implements a series of matched filters at a number of orientations on each frame of the SMLM acquisition [4, 5]. Using a finite number of these candidate filters leads to some orientations being more efficiently detected than others and errors in initial orientation estimates, which can become computationally expensive to avoid by increasing the number of oriented filters used. Deep-learning approaches to PSF detection often use even more convolutions, typically in the range of hundreds to thousands [6, 7]. Here, we use a steerable filter design and an edge orientation algorithm developed by Freeman and Adelson [8] to detect DH PSFs and analytically estimate their orientation with just 7 (separable) convolutions. This detection algorithm can be automatically tuned to a given DH PSF, and is normalized to be compatible with signal-to-noise-based thresholding [9]. Combined with the highly parallelized localization analysis architecture of the PYthon Microscopy Environment (PYME), our plug-in enables a high-performance pipeline for DH PSF SMLM at state-of-the-art quality, without manual tuning.

## Method

Optimal steerable filters have been developed for several SMLM use cases [10], including DH PSF detection [11]. Here, we instead implemented the 2^nd^ derivative of a two-dimensional Gaussian filter (*G*_2_) because it has a low number of basis filters and there exists an analytic expression for the orientation that maximizes the response at each pixel as previously derived [8]. We normalize the 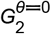 filter as

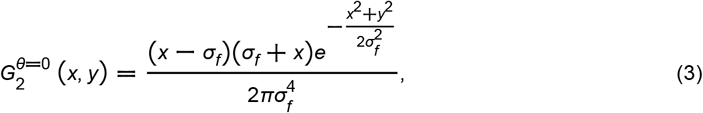

where *σ*_*f*_ is the filter width parameter (See S1.1). *gg*(*y* = 0),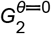 (*y* = 0), and their squared convolution are shown in Figure 1E. Despite the discrepancy between the filter and the model, the squared filter output still peaks at the center of the model as desired, provided *σ*_*f*_ is chosen within a reasonable range. After convolving a raw camera frame with just 3 oriented 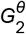 filters, a linear combination of these 3 convolutions can be used to construct the frame convolved with any orientation of the *G*_2_ filter without performing the final desired convolution explicitly [8]. Further, there exists a set of separable basis kernels (as shown in figure 1F).

The Hilbert transform of *G*_2_, *H*_2_, can be reasonably approximated with a 3^rd^-order polynomial, and requires 4 basis filters to steer [8] (see also [13]). These Hilbert transformed filters (Fig. 1F) share the same spatial frequency response, but with a phase shift from the *G*_2_ filtering [14]. Freeman and Adelson show that adding the *G*_2_ and *H*_2_ filtered outputs in quadrature simplifies to a Fourier series in *θ*, from which the strength of the response and the angle maximizing it are well approximated using the lowest frequency term [8]. This procedure results in filtered strength (Fig. 1G) and orientation maps (Fig. 1H) of the input image.

To detect candidate molecules across a large dynamic range of brightness and background with minimal user input, we use percentile-based background-subtraction and per-pixel signal-to-noise (SNR)–based thresholding [9], both of which are already implemented in PYME. A noise map, *σ*_noise_, is calculated on each frame as the expected standard deviation given shot-noise, excess noise (for EMCCDs [15]) and read-noise (permitting use of calibrated camera maps [9]). Multiplying the noise map by a threshold factor, *T*, allows for SNR-based detection on normalized filter output of each frame. To instead perform SNR-based thresholding on the quadrature strength map, a threshold map is computed as *c* (*σ*_noise_*T*) ^2^ where *c* is an analytic normalization factor accounting for the center-pixel filter response to the double Gaussian function of eq. 2 (see section S1.1). The strength map and threshold map are each maximum filtered in order to extract candidate molecule positions at strength map maxima which correspond to DH PSF images with lobe brightnesses sufficiently above noise (see section S1.3). A precise orientation estimate for each candidate can then be extracted from the orientation map (Fig. 1H). High-quality DH PSF detections and orientation estimates are therefore extracted from each SMLM frame with just 7 convolutions and 2 maximum filters (Fig. 1I).

Fitting of candidate molecules is performed using equation 2 as the model function, with an analytic Jacobian, and a least squares optimization (through SciPy [16] scipy.optimize.leastsq) with residuals weighted by 1/*σ*_noise_. To preserve the noise model in the weights, the per-pixel background estimate is added onto the model function during evaluation (rather than subtracted from the data). Fitting each frame of a DH PSF Z-stack calibrates the orientation (*θ*), lateral wobble, and other DH PSF parameters as a function of axial defocus. After localizing a SMLM dataset, a spline interpolation of the calibrated orientation *θ*_cal_ (*z*) is used to look-up the z position for each localization. Lateral wobble is similarly corrected with a spline interpolation. The uncertainty estimate in *θ* from fitting each localization, *δθ*_*i*_, is propagated to an uncertainty in *z*_*i*_ using the slope of *θ* (*z*_*i*_), and localizations are additionally annotated with residuals from other calibrated values, e.g. *L*_*i*_ − *L*_cal_ (*z*_*i*_), to facilitate downstream quality control on localizations.

## Results

Combining our effectively tuning-free detection with the strengths of an analytic model function makes achieving state-of-the-art localization quality remarkably straightforward. The single filter hyperparameter, *σ*_*f*_ is automatically determined during DH PSF calibration (see Figure 2A,B and section S1.2), as are the DH PSF parameters used as initial guesses for fitting (Fig. 2C). As the SNR-based thresholding obviates the need to tune a threshold, the entire detection can proceed successfully with no manual tuning (Fig. 2D).

**Figure 2:**
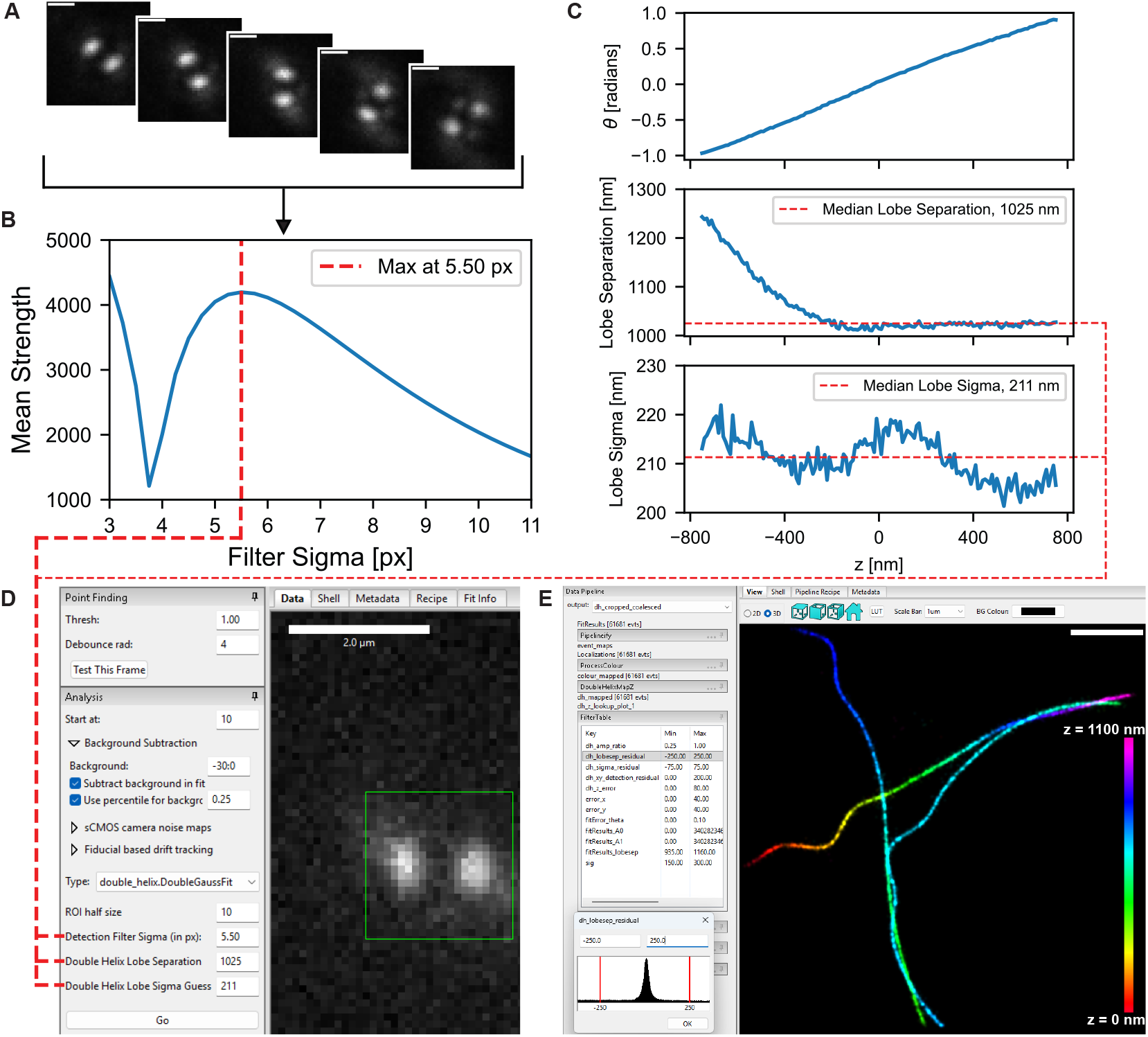
A) Slices from a calibration bead z-stack from reference [12] corresponding to the z positions −800 nm, −400 nm, 0 nm, 400 nm, 800 nm. Scale Bar: 1 µm B) Output of the Optimize DH PSF Detection function. Mean detection strength at the DH PSF center pixel for each slice shown in (A) is shown as a function of filter sigma. The function identifies the filter sigma that peaks the detection strength. C) Output of the Calibrate DH PSF function. *θ*, lobe separation, and lobe sigma of the calibration bead is plotted as a function of z. Plots of lobe separation and lobe sigma indicate the median lobe separation and median lobe sigma across the z range for use as initial guesses for DH PSF fitting. D) Screen capture of the Point Finding and Analysis modules in PYMEImage along with the MT0.N1.LD-DH dataset. Thresholding with a value of 1.00 translates roughly to thresholding with an SNR of 1. The optimal filter sigma identified in (B) and the median lobe separation and median lobe sigma identified in (C) are used as analysis parameters. The green box indicates a detected DH PSF resulting from the Test This Frame function. Scale Bar: 2 µm E) Screen capture of the MT0.N1.LD-DH dataset rendered in PYMEVisualize. The left-hand side shows the available filtering parameters with a histogram of dh_lobesep_residual shown below as an example. The red bars in the histogram indicate the lower and upper cutoffs for filtering. Localizations are rendered with 30-nm pointsprites with an alpha of 0.055 and colored by *z* with a range of *z* =0 nm to 1100 nm.

Using an analytic model and a structured PSF delivers abundant quality metrics to filter out spurious or poor-quality localizations. These metrics include typical uncertainties, sizes of PSF features, deviation from calibrated PSF feature size based on estimated *z* position, and deviation of final fitted lateral position from initially detected position, and are shown in table 1. These metrics can be leveraged to easily perform adequate filtering, by filtering with especially liberal bounds on all parameters (e.g. localization uncertainty in *x* and *y* < 40 nm, and in *Z* < 80 nm. See also table 1). In this way, manual tuning is also not typically required to achieve quality controlled localizations (Fig. 2E).

**Table 1:**
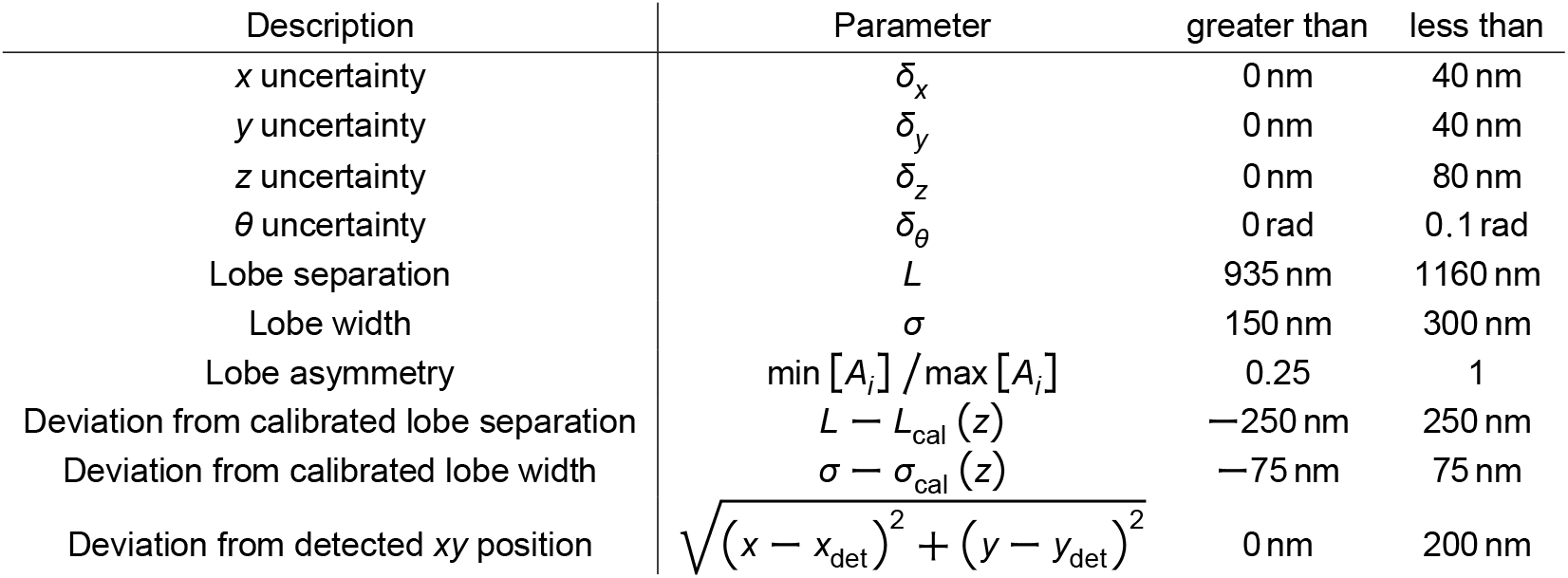
Localization filter parameters used after localizing the MT0.N1.LD-DH dataset.

| Description | Parameter | greater than | less than |
| --- | --- | --- | --- |
| x uncertainty | $\delta_x$ | 0 nm | 40 nm |
| y uncertainty | $\delta_y$ | 0 nm | 40 nm |
| z uncertainty | $\delta_z$ | 0 nm | 80 nm |
| $\theta$ uncertainty | $\delta_\theta$ | 0 rad | 0.1 rad |
| Lobe separation | $L$ | 935 nm | 1160 nm |
| Lobe width | $\sigma$ | 150 nm | 300 nm |
| Lobe asymmetry | $\min[A_i] / \max[A_i]$ | 0.25 | 1 |
| Deviation from calibrated lobe separation | $L - L_{\text{cal}}(z)$ | -250 nm | 250 nm |
| Deviation from calibrated lobe width | $\sigma - \sigma_{\text{cal}}(z)$ | -75 nm | 75 nm |
| Deviation from detected xy position | $\sqrt{(x - x_{\text{det}})^2 + (y - y_{\text{det}})^2}$ | 0 nm | 200 nm |

We benchmark this performance by localizing the MT0.N1.LD-DH dataset, a synthetic training dataset from the 3D SMLM software fight-club publication [12] (Fig. 3A). Our detection parameters were determined automatically from the calibration procedure as shown in figure 2, and a detection threshold of 1.0 was used. Relaxed quality filtering was performed using the bounds shown in table 1. Comparing the resulting localizations with the ground truth dataset (for details, see S3), our pipeline achieves a Jaccard index of 82.7%, lateral root mean squared error (RMSE) of 16.1 nm, and axial RMSE of 21.4 nm. On a comparable Sage et al. dataset which is not publicly available, MT1.N1.LD-DH, the top performing software was SMAP-2018 [17] with a Jaccard Index of 77.1%, Lateral RMSE of 27 nm, and Axial RMSE of 20.9 nm. Our results compare favorably (see table 2).

**Table 2:**
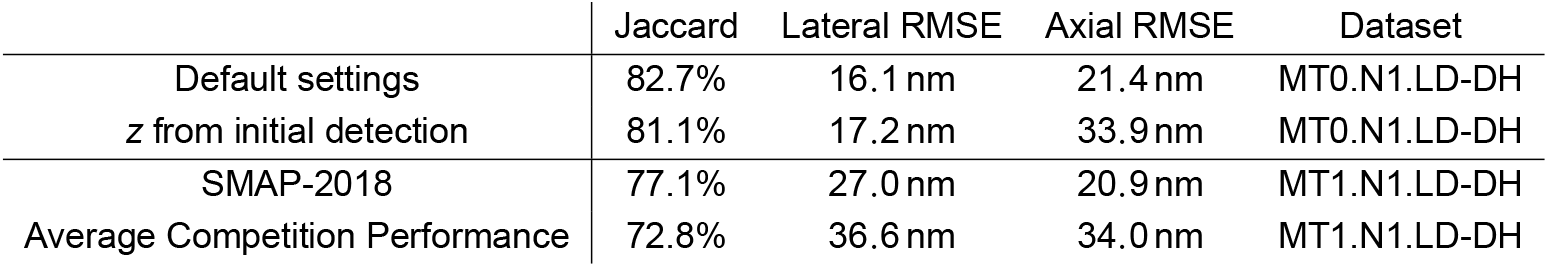
Performance comparisons using high SNR, low density synthetic datasets from Sage et al. [12].

**Figure 3:**
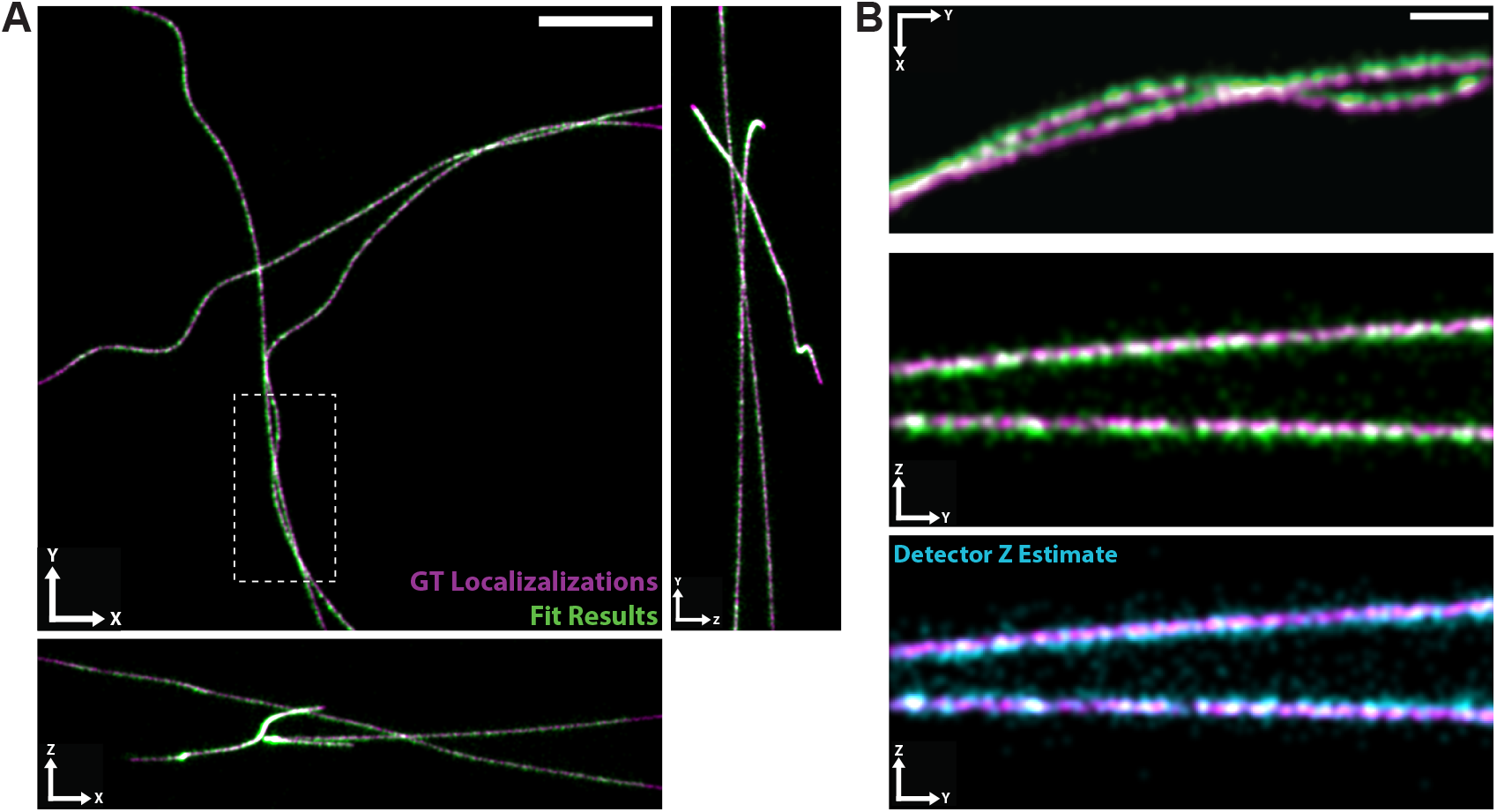
A) Ground truth positions (magenta) and localization fit results (green) for the full field of view of the MT0.N1.LD-DH dataset shown in *x*-*y, x*-*z*, and *y*-*z* projections. White dashed box indicates ROI shown in (B). Scale Bar: 1 µm B) Ground truth positions (magenta) and localization fit results (green) for the region in (A) indicated by the white dashed box. Localizations are shown in *x*-*y* (top) and *y*-*z* projections (middle). Bottom row shows ground truth positions (magenta) and localizations with *z* estimated from the detector *θ* (cyan) for the same ROI from a *y*-*z* projection. Scale Bar: 500 nm.

We note that the most obvious discrepancies between our localizations and the ground truth molecules are a lateral offset potentially introduced by imperfect wobble correction (at expense to the Lateral RMSE) and missing detections on the lateral border (at expense to the Jaccard index). While the performance scoring removes 450 nm lateral borders to allow comparison with results from reference [12], the lobe separation of *L* = 1025 nm means the scoring includes penalties for ground truth molecules whose DH PSF image may be cropped to essentially a single lobe. Missing detections at the border are also to be expected due to our filtering kernel size of 21 × 21 pixels (2.1 µm × 2.1 µm), however this is easily remedied in practice by imaging more generous fields of view.

To characterize whether the detection threshold was indeed unnecessary to tune manually, we swept the threshold factor, *T*, from 0.5 to 2.0. (see Figure S1). We found that the filtering of table 1 ensured very reasonable performance even for low SNR thresholds, while the quality benefit to only detecting at higher SNRs of 2*σ*_noise_ was minimal as one expects due to the square root relationship typical of localization precision and brightness.

When run on a single Window’s 11 PC with a 3.4 GHz Intel i7-14700K CPU and 32 GB of RAM, the 19,996 frame MT0.N1.LD-DH dataset is localized in 71.6 s including all file I/O. Profiling a computation thread during this benchmark using PYME’s mProfile shows that the fitting of candidate molecules takes about 30 times longer than the detection. The speed of a 2D convolution using a separable *k*×*k* filter is expected to improve by *k*^2^/(2*k*) when performed as 2 × 1D convolutions using k-length kernels. With *k* = 21, on the first frame of the 64 × 64 pixel MT0.N1.LD-DH dataset, executing the 7 detection filters and calculating the orientation and strength maps decreases in runtime by a factor of 37.5 using separable filters, surpassing the expected factor of 10.5 (presumably due to implementation details in SciPy). Notably, if higher-throughput is required, the runtime can be readily decreased by enlisting more computers into the PYME cluster [18].

To benchmark the feasibility of using steerable filters in place of fitting for low-latency applications such as focus stabilization or trapping, we substituted the initially detected *θ* value for the fitted *θ* value when determining the *z* position for each localization, leaving the other filters in place (cyan localizations in Fig. 3B). After removing a constant (median) offset in *z*, our detection-only *z* result still maintains reasonable quality, with an axial RMSE typical of average performance in the original 3D software challenge [12] (see tables 2, S1 and Fig. S2). This average performance is in spite of the *z* estimates near the lateral borders breaking down significantly (see Fig. S2). Thus, the detector’s orientation estimate provides a feasible means to low-latency *z* estimation with adequate precision for several applications.

## Discussion

While localization analysis is by now a fairly mature field, detecting and fitting engineered PSFs pose additional challenges which are often less tractable analytically and pose additional computational burdens. This has pushed many towards deep-learning-based approaches, which are quite powerful in their own rite, however, analytic models can have advantages in terms of speed, transferability, robustness, and explicit nature. Here, we repurposed a steerable filter set to accomplish DH PSF detection in a computationally efficient manner without orientation sub-sampling or resulting bias. We deployed this detection and accompanying double Gaussian fit as a plug-in for the PYthon Microsocpy Environment (PYME) in order to leverage its SNR-based thresholding, percentile-based background estimation, and performant (distributed) analysis capabilities [18, 19]. Combined with a quick calibration routine, we demonstrated that this pipeline enables high-quality SMLM reconstruction from DH PSF data with minimal user input.

We believe this software is immediately useful to users of DH PSFs. In addition to full SMLM analysis, we highlight that our detection algorithm alone provides reasonable quality super-resolved *z* localization, which may benefit applications such as real-time drift correction or trapping where iterative optimization can be prohibitive. We expect steerable filters to continue to find use in single-molecule image analysis for complex PSFs, ranging from fixed dipole emitters imaged with open-aperture PSFs [10] to engineered PSFs including the double helix. Orientation estimation from steerable filters might be developed for filter sets which are more optimally matched to DH or other engineered PSFs.

## Acknowledgment

We are especially grateful to David Baddeley for developing and maintaining PYME. We appreciate useful discussions with the lab of Anna-Karin Gustavsson. W.E.M. is a Sarafan ChEM-H Institute Fellow.

## Data availability

The described software is available at reference [20]. The simulated DH PSF SMLM data is the MT0.N1.LD-DH dataset from reference [12] obtained from srm.epfl.ch/Challenge/ChallengeSimulatedData. The remaining data and code are available from reference [13].

