## Supplementary Materials for "Efficient Double Helix Detection with Steerable Filters"

### Supplementary Materials
### Efficient Double Helix Detection with Steerable Filters


Andrew E. S. Barentine [1,*,✉], Ashwin Balaji[1,2,*], and W.E. Moerner [1,3]

[1]Department of Chemistry, Stanford University, Stanford, California 94305, USA

[2]Biophysics PhD Program, Stanford University, Stanford, California 94305, USA

[3]Sarafan ChEM-H, Stanford University, Stanford, CA 94305, USA

*these authors contributed equally

✉


April 3, 2026

#### S1 Detection

##### S1.1 Filter construction and normalization

Freeman and Adelson developed a general image analysis algorithm to estimate local orientation using their $G_2$ and $H_2$ quadrature steerable filter set [1]. Their $G_2^{\theta=0}(x, y)$ filter is given as

$$0.9213(2x^2 - 1)e^{-(x^2+y^2)}, \tag{S1}$$

which does not contain a width parameter necessary in our application to scale the filter to detect different DH PSFs. We instead built our $G_2$ filter set by differentiating a sum-normalized Gaussian, arriving at

$$G_{2a}(x, y) = G_2^{\theta=0}(x, y) = \frac{(x - \sigma_f)(\sigma_f + x)e^{-\frac{x^2+y^2}{2\sigma_f^2}}}{2\pi\sigma_f^4}, \tag{S2}$$

which contains the width parameter, $\sigma_f$. While $G_2$ filters themselves integrate to zero, integrals of only the negative or positive regions both scale with $1/\sigma_f^2$, which complicates the optimization of $\sigma_f$ for matched filtering of a given DH PSF. Our $G_{2a}$ has therefore been multiplied through by an additional factor of $\sigma_f^2$ to remove this dependence from $G_2$ filtered outputs. We used our $G_{2a}$ to derive a polynomial approximation of its Hilbert transform, $H_{2a}$, and then derived expressions for the separable basis functions for the $G_2$ and $H_2$ steerable filter sets following the procedure established by Freeman and Adelson (See [2]).

To perform SNR-based thresholding on the strength map, $S$, resulting from quadrature filtering, we scale the noise map by the center-pixel response, $S_o$, evoked by the double Gaussian function of eq. 2, which is given by:

$$S_o = \frac{\pi A^2 \sigma^4 \sigma_f^4}{(\sigma^2+\sigma_f^2)^3} \Bigg[ \operatorname{erf}\left(\frac{1}{2\sqrt{2}\sqrt{\sigma^2+\sigma_f^2}}\right)^2 e^{-\frac{(L+1)^2}{4(\sigma^2+\sigma_f^2)}} \left((L-1)\left(-e^{\frac{L}{2(\sigma^2+\sigma_f^2)}}\right)+L+1\right)^2$$

$$- e^{-\frac{L}{4(\sigma^2+\sigma_f^2)}} \left(\operatorname{erf}\left(\frac{L-1}{2\sqrt{2}\sqrt{\sigma^2+\sigma_f^2}}\right) - \operatorname{erf}\left(\frac{L+1}{2\sqrt{2}\sqrt{\sigma^2+\sigma_f^2}}\right)\right)^2 \Bigg]^{1/2} \quad \text{(S3)}$$

where erf is the Gauss error function and the amplitudes of the double Gaussian function are assumed to be equal: $A = A_1 = A_2$. For a given DH PSF and filter set, $S_o$ reduces to $c \cdot A^2$, where c is a constant (see [2]). The initial guesses for $L$ and $\sigma$ are used to identify the value of $\sigma_f$ that peaks the analytical center-pixel detection strength. This $\sigma_f$, along with the initial guesses for $L$ and $\sigma$ are then used to compute $c$. A scaled threshold map is finally computed as $c\left(\sigma_{\text{noise}} T\right)^2$ which allows SNR-based thresholding on the strength map.

#### S1.2 Filter optimization

Detection benefits from tuning the filter width, via the $\sigma_f$ parameter, to match a given DH PSF. An optimized $\sigma_f$ for detection, and initial guesses for $L$ and $\sigma$ for fitting are extracted from a z-stack of a single, centered calibration object such as a sub-diffraction fluorescent bead (Fig. 2A). While PYME has functions for PSF extraction averaging over multiple beads and deconvolving the bead shape itself, here we simply apply a lateral crop to a z-stack of beads. A fit of the DH PSF model function is then performed on each slice. Slices with fitted $\theta$ displacements of approximately $-80$, $-40$, $0$, $40$, and $80$ degrees relative to the central z slice are used to calculate an average center-pixel strength for a given test $\sigma_f$ value (Fig. 2B). Test $\sigma_f$ values to maximize this average range from the estimated $\sigma$ to the estimated $L$. The lower bound is especially important because at $\sigma_f$ smaller than the DH PSF lobe $\sigma$, high spatial frequency patterns are produced without a peak at the DH PSF center.

#### S1.3 Candidate extraction

With an optimized $\sigma_f$, the strength image maxima occur at DH PSF centers, $\left(x_o, y_o\right)$, while the scaled threshold map one would like to compare it to instead peaks at the DH PSF lobes. We therefore perform a maximum filter on both the strength image, and the threshold-scaled noise map before comparison, both using kernel sizes of $\lceil L+\sigma \rceil$ pixel $\times \lceil L+\sigma \rceil$ pixel where $\lceil\rceil$ denotes a ceiling operation. The detection thresholding is then finally performed with binary operations, constructing a boolean map of candidate molecule positions as

$$(\text{MaxFilter}[S] == S) \text{ AND } \left(S \geq \text{MaxFilter}\left[c\left(\sigma_{\text{noise}} T\right)^2\right]\right). \quad \text{(S4)}$$

The row and column positions where this evaluates to True are taken to be candidate molecule positions for subsequent fitting, and these pixelated positions are used to extract initial $\theta$ guesses from the orientation map.

#### S2 DH PSF Calibration and post localization mapping

Calibration of the DH PSF is necessary to produce a mapping from a localization's fitted $\theta$ to its localized $z$ position. A PSF z-stack is fit with the model function on each slice, producing a calibration file which records the z dependence of the following parameters: $\theta$, $L$, $\sigma$, $x$, and $y$. The $x$ and $y$ variation is referred to as lateral wobble. Calibration also plots $\theta$, $L$, and $\sigma$ as a function of $z$ for a user. Additionally, the median values of $L$ and $\sigma$ are identified for use in detection normalization and as initial guesses for fitting. Thus, the optimization and calibration procedures provide users with parameter values for localization with minimal user input (Fig. 2C, D). However, it is recommended to test detection with these parameters on the calibration z stack to ensure that the identified parameters effectively concentrate the detection strength in the center of the DH PSF across the desired z range. We find that small increases in $\sigma_f$ ($\sim$0.25 - 0.5 px) may be helpful in slightly improving the detection at the end range of some DH PSF that exhibit more change in $L$ over their axial range.

After localization fitting, a cubic spline is used to interpolate between points in the $\theta$ vs $z$ calibration and map localizations in $z$. The lateral wobble vs $z$ is similarly corrected with a cubic spline interpolation of the calibration. Localizations can then be filtered on several DH PSF parameters to ensure the final localizations include reasonable single-molecule fits (Fig. 2E). Filtering parameters include $x$ uncertainty, $y$ uncertainty, $z$ uncertainty, $\theta$ uncertainty, lobe separation, lobe width, lobe brightness asymmetry, deviation from calibrated lobe position, deviation from calibrated lobe width, and deviation from detected $x, y$ position (see Table 1).

#### S3 Performance benchmarking on ground truth synthetic data

Comparison with the ground truth dataset required swapping $x$ and $y$, shifting $x$ and $y$ by 1 pixel (100 nm) and shifting $z$ by 750 nm (half the simulated $z$ range). Following the procedure outlined in Sage et al [3], we cropped the 6.4 μm $\times$ 6.4 μm field of view by 450 nm on each lateral border, and linked localizations to their nearest neighbor ground truth molecule within 250 nm. The resulting Jaccard index of 82.7%, lateral root mean squared error (RMSE) of 16.1 nm, and axial RMSE of 21.4 nm compares favorably with the best in class software from the challenge (on a comparable Sage et al dataset which is not publicly available, MT1.N1.LD-DH, the top performing software was SMAP-2018 with a Jaccard Index of 77.1%, Lateral RMSE of 27 nm, and Axial RMSE of 20.9 nm).

##### S3.1 Detection threshold effect on final outcome

We tested the effect of varying the signal-to-noise (SNR) based detection thresholding on the final comparison by sweeping the threshold factor from 0.5 to 2.0. All other localization analysis parameters were kept constant, including post-localization filtering with parameters shown in Table 1. As expected, restraining the detection to higher SNR detections serves to increase the axial and lateral RMSEs at some expense to the Jaccard index (see figure S1), primarily due to a decline in Recall (number of true positive detections divided by the number of ground truth positions).

##### S3.2 Detection-only Z performance

This detection-only $z$ performance decreases near the borders. We highlight several examples in Figure S2. The rendered images do not show detections or ground truth molecules within 450 nm of lateral borders following the procedure in reference [3]. The median lobe separation of $L = 1025$ nm means that some orientations of DH PSFs near the borders might be cropped to essentially a single lobe, which would affect detection. The inaccurate detection-only $z$ values for emitters which are near the borders is likely due to our filter kernel size of 21 x 21 pixels (2.1 μm $\times$ 2.1 μm). We note that excluding an additional 500 nm of the

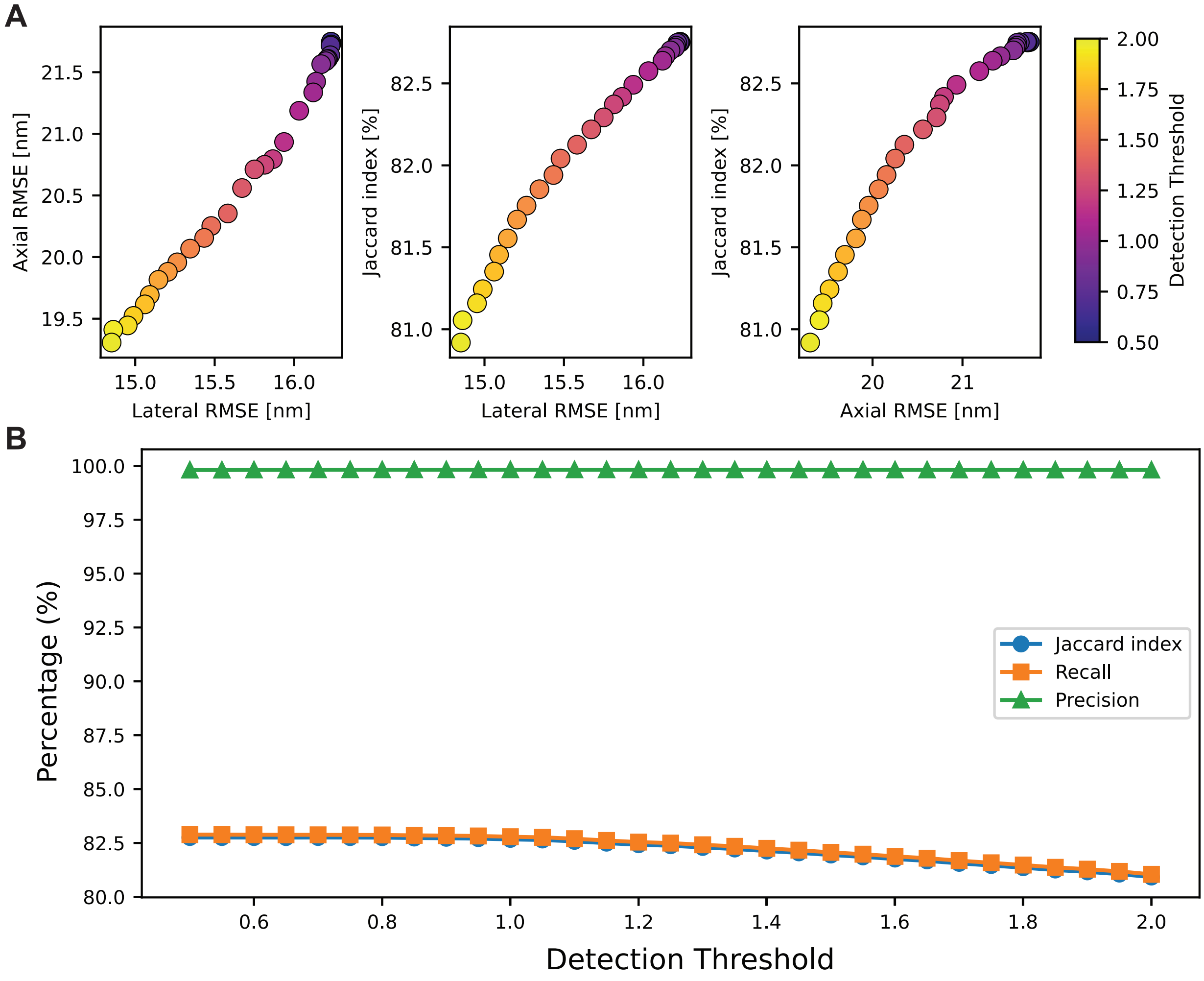


Figure S1: Dependency of Benchmarking Metrics for the MT0.N1.LD-DH Dataset on Detection Threshold: A) Left to right: Axial RMSE vs Lateral RMSE, Jaccard Index vs Lateral RMSE, and Jaccard Index vs Axial RMSE colored by detection threshold. Detection threshold was varied from 0.5 to 2.0 in increments of 0.05. B) Jaccard Index, Recall, and Precision (as defined in Sage et al [3]) as a function of detection threshold.

border improves the detection-only *z* Jaccard Index by 1.7% due to a minor change in linking to ground truth molecules, but also improves the axial RMSE by 5.7 nm (see *z from initial detection, without border* entry in table S1).

| | Jaccard | Lateral RMSE | Axial RMSE | Dataset |
|---|---|---|---|---|
| Default settings | 82.7% | 16.1 nm | 21.4 nm | MT0.N1.LD-DH |
| *z* from initial detection | 81.1% | 17.2 nm | 33.9 nm | MT0.N1.LD-DH |
| *z* from initial detection, without border | 82.8% | 16.8 nm | 28.2 nm | MT0.N1.LD-DH |
| SMAP-2018 | 77.1% | 27.0 nm | 20.9 nm | MT1.N1.LD-DH |
| Average Competition Performance | 72.8% | 36.6 nm | 34.0 nm | MT1.N1.LD-DH |

Table S1: Performance comparisons using high SNR, low density synthetic datasets from Sage et al [3].

### S4 Rendering

The image shown in Figure 2E was displayed in PYMEVisualize [4] using 30-nm pointsprites with an alpha of 0.055. Localizations in Figure 3 and Figure S2 were rendered in PYMEVisualize [4] as isotropic 3D Gaussians with $\sigma = 10$ nm into 3D image stacks that were pixelated with isotropic 5 nm pixels. Final images of *x*-*y*, *x*-*z*, and *y*-*z* projections were then rendered as mean projections of these stacks. See [2] for the full PYME recipes.

A

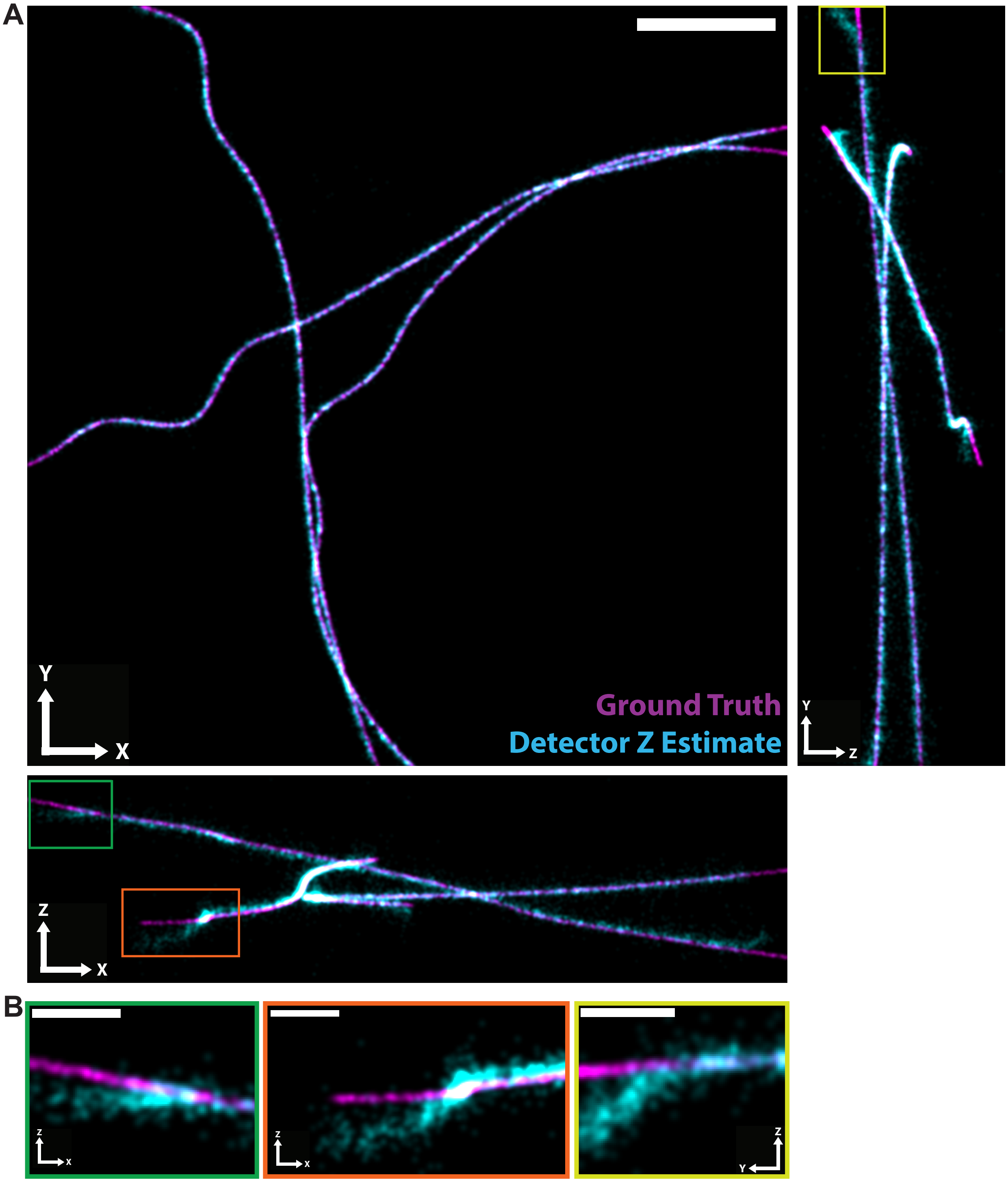


Figure S2: A) Ground truth positions (magenta) and localizations with *z* estimated from the detected *θ* (cyan) for the full field of view of the MT0.N1.LD-DH dataset shown in *x*-*y*, *x*-*z*, and *y*-*z* projections. Colored boxes (green, orange, yellow) indicate ROIs shown in (B). Scale Bar: 1 µm. B) ROIs indicated by green, orange, and yellow boxes in (A) shown from *x*-*z* (green, orange) or *y*-*z* (yellow) projections. Localizations with *z* estimated by detected *θ* (cyan) show deviation from ground truth positions (magenta) for these ROIs close to the edge of the field of view. Scale Bars: 200 nm.
